# Tracing the Evolutionary Origin of Interferon to Basal Chordates and Unveiling Its Antiviral Functionality in Jawless Vertebrates

**DOI:** 10.64898/2026.05.30.728932

**Authors:** Xinli Wang, Qinghuan Wang, Anni Xie, Lisi Deng, Ziwen Huang, Yiming Zhao, Yange Cao, Rong Fu, Weicong Huo, Yanhong Chen, Guang Li, Anlong Xu, Shaochun Yuan

## Abstract

Interferons (IFNs) are essential mediators of antiviral defense in vertebrates, having gradually replaced the RNA interference (RNAi) antiviral mechanism that predominates in invertebrates and plants. To date, IFNs have been identified exclusively in jawed vertebrates, leaving the origin of IFN-based antiviral mechanisms largely mysterious. In this study, by conducting a genome-wide screening of IFN homologs accross various species, we successfully identified seveal IFN homologs from agnatha and lancelet, but not other invertebrates. Notably, both agnatha and lancelet IFN homologs have ability to induce a set of interferon-stimulated genes (ISGs)-like genes, thus tracing the origin of IFN to basal chordate. Using VSV infected peripheral blood mononuclear cells (PBMCs) of Japanese lampreys, we found that lamprey IFNs have antiviral functionality by inducing the expression of hundreds of ISGs through interacting with a heterodimeric complex composed of CRFB7 and CRFB14. In addition to robustly mediating antiviral responses in monocytes, lamprey IFNs exert their effects on variable lymphocyte receptor B (VLRB)^+^ cells by remodeling the cytokine/chemokine networks to orchestrate antiviral innate and adaptive immunity. Furthermore, cross-species functional comparison of Dicer revealed that changes in residues essential for dsRNA processing occurred concurrently with the evolution of the IFN system. Collectively, these findings uncover the evolutionary origin of IFN and underscore its ancient roles in antiviral response and immune regulation, especially in the takeover of the RNAi antiviral mechanism during the early evolution of vertebrates.

**Significance Statement:** Interferons (IFNs) represent a hallmark of vertebrate antiviral immunity, yet their origin has remained elusive. Here, we reported bona fide IFN ligands and cognate receptors in both lamprey and lancelet, two key species placed at the transition from invertebrates to vertebrates. Using the VSV infection model, we further demonstrated the roles of lamprey IFN in antiviral defense and immune regulation. We also found that substitutions in residues essential for Dicer mediated dsRNA processing coincided with the emergence of the IFN system. Collectively, our findings provide key insights into the evolutionary origin of IFN-based immunity and its gradual replacement of RNAi as the dominant antiviral strategy in vertebrates.

## Introduction

IFN is a class of small and inducible glycoproteins with diverse biological activities. It was first discovered in the 1950s by Alick Isaacs and Jean Lindenmann (1), who observed its ability to “interfere” with viral replication within cells, thereby earning its name. In the decades following its discovery, four types of IFN were identified in jawed vertebrates based on their structure and activities, including type I, II, III, and IV (2). Type-I IFN (IFN-I) has been widely recognized as an essential mediator in triggering antiviral immunity in jawed vertebrates. In mammals, upon viral infection, host cells detect viral nucleic acids via pattern recognition receptors (PRRs) such as RIG-I-like receptors (RLRs), Toll-like receptors (TLRs), and cyclic GMP-AMP synthase (cGAS) (2–5). These receptors trigger downstream signaling pathways involving adaptor proteins like MAVS (Mitochondrial antiviral signaling protein) and STING (Stimulator of interferon genes), leading to the production of IFN-Is and other antiviral cytokines. After being released by viral infected cells, IFN-I binds to its receptor, a heterodimeric complex composed of IFNAR1 and IFNAR2, to activate the JAK (Janus kinase) -STAT (Signal transducer and activator of transcription) signaling and induce the production of ISGs (Interferon stimulated genes) that play critical roles in mediating antiviral responses and immune regulation (2, 6, 7).

While IFNs are central to antiviral responses in jawed vertebrates, plants and invertebrates like *Drosophila* and nematodes applied RNAi, an evolutionarily conserved mechanism of gene silencing in eukaryotes, as a pivotal antiviral defense mechanism (8). In the antiviral process mediated by RNAi, viral double-stranded RNA (dsRNA) is recognized and processed by the enzyme Dicer into small interfering RNAs (siRNAs). These siRNAs then guide the RNA-induced silencing complex (RISC) to degrade viral RNA transcripts with complementary sequences, thereby inhibiting viral replication and spread within host cells (8). Although RNAi robustly protects hosts from viral infection, its natural antiviral function in vertebrates is limited, such as zebrafish, mouse and humans (8, 9). In 2013, using Nodamura virus (NoV) and its mutant (NoVΔB2) that lacks the B2 protein, a viral suppressor of RNAi (VSRs), the antiviral function of RNAi in mammals is evidenced (10). After that, studies have gradually unveiled other VSRs that directly inhibit the host RNAi pathway, such as the NS1 protein of influenza virus and the NS3 protein of dengue virus, which can prevent the cleavage of viral dsRNA by host Dicer (11). Addition to VSRs, the IFN system in mammalian cells can also inhibit the RNAi-mediated antiviral functions. For example, LGP2 (Laboratory of genetics and physiology 2), one of the ISGs, can competitively bind to dsRNA with Dicer, thereby suppress the ability of Dicer (12). In stem cells that lack the IFN antiviral response, an isoform of Dicer named antiviral Dicer (aviD) was identified to efficiently cleave viral dsRNA to orchestrate antiviral RNAi, thereby protecting mammalian stem cells from RNA virus infection, including Zika virus and SARS-CoV-2 (13). These mechanisms highlight the evolutionary “takeover” of RNAi by the IFN system, particularly in differentiated cells. Thus, the emergence of IFN in evolution represents a milestone in this transition.

Potentially due to the intense pressure of viral infection, IFN evolved rapidly by gene expansion and functional diversification across species, especially for IFN-I (14, 15). One of the notable events is the emergence of intronless IFN-I genes in amphibians, which bridges the evolutionary gap between aquatic and terrestrial vertebrates (16). These intronless IFN genes are thought to have arisen through retrotransposition events, and subsequently expanded and diversified during evolution (17). For example, IFN-α has undergone significant expansion in humans, mice, and bats (2, 18). Humans possess at least 13 subtypes of IFN-α which are all located on chromosome 9 and share high sequence homolog (3). IFN-τ is specifically presented in ruminants and associated with pregnancy maintenance, while IFN-δ is only found in pigs and likely related to antiviral immunity and immune regulation (19, 20). While mammals exhibit an expanded IFN-I repertoire, some teleost fish display an even more dramatic expansion of this gene family, with rainbow trout (*Oncorhynchus mykiss*) and Atlantic salmon (*Salmo salar*) each encoding dozens of IFN-I genes (21, 22). Notably, all IFN-Is in teleost fish are intron-containing IFN-I genes which can be further subdivided into distinct groups based on the number of cysteine bridges and their phylogenetic relationships (22). For instance, zebrafish (*Danio rerio*) encodes four IFN-Is belong to group I (IFNphi1 and IFNphi4) and group II (IFNphi2 and IFNphi3) (21, 22). Another notable event is the absence of IFN-III and the segmental duplication of IFN-II in teleost fish, giving rise to the IFN-γ-related gene (IFN-γrel) in fish (23). This fish-specific paralog plays crucial roles in immune regulation and antiviral responses (24).

Although extensive efforts have been dedicated to identify IFNs across various species, they have been exclusively found in jawed vertebrates based on sequence identity according to current literatures (14, 15). Vago, a molecule characterized by a single von Willebrand factor C domain (SVWC), has been identified as an antiviral cytokine-like molecule in arthropods (25–27). However, it is not a member of IFN family. Thus, the origin of IFNs and the process by which they gradually replaced RNAi in antiviral responses among ancient species remain unclear. In this study, we successfully identify IFN homologs and their heterodimeric receptor complex in jawless vertebrates and lancelets by employing high-throughput screening in jawless and invertebrates, thus tracing the origin of IFN to basal chordates. Sharing a common ancestor with jawed vertebrates, lamprey IFNs not only play a crucial role in antiviral responses by robustly upregulating a suite of ISGs, but also participate in orchestrating the activation of VLRB^+^ cells and monocytes, highlighting its multifaceted functions in immune regulation. Notably, comparative analyses revealed the functionally consequential amino acid substitutions of Dicer that coincided with the emergence of the IFN-mediated antiviral system, offering new insights into the functional shift from RNAi-based to IFN-based antiviral defense in early vertebrate evolution.

## Results

### Discovery of IFN homologs in agnatha and lancelet, tracing the evolutionary origin of IFN to basal chordates

IFN is a group of type II α-helical cytokines that exhibits some classical characteristics. First, it is a protein with approximately 200 amino acids, featuring a 22-23 amino acid N-terminal signal peptide. Second, the protein consists of five α-helices and is encoded by five exons. Third, the classical IFN is proposed to have a 0-0-0-0 exon-intron phase in the genomic coding sequence (28) **(Fig. 1A)**. To discover whether there are IFN homologs in jawless vertebrates and invertebrates, we developed a script to screen genes with 0-0-0-0 intron-exon phase of the entire genome. We first searched the genome of sea lamprey (*Petromyzon marinus*, Pma), and finally obtained 132 genes. However, only 14 genes contained a signal peptide and had protein lengths of less than 300 amino acids. Then, ESMFold was used to find whether these 14 proteins displayed secondary α-helix structures, and only one molecule (*LOC116947453*, named as *PmaIfn1*) was finally filtered out. Since IFNs may be clustered in the same chromosome, we further screened genes near *LOC116947453* and identified two genes in sea lamprey on the same chromosome, which were named *PmaIfn2* and *PmaIfn3* **(listed in SI Appendix, Dataset S1)**. Unlike *PmaIfn1*, *PmaIfn2* and *PmaIfn3* were encoded by four exons with intron phases of 0-0-0 due to the fusion of exons 4 and 5 (**Fig. 1B and SI Appendix, Dataset S1**). Using the same screening pipeline, we further identified *EA37267* (*EatIfn*) from hagfish *Eptatretus atami*, three molecules (*LreIfn1* - *LreIfn3*) from lamprey *Lethenteron reissneri* (Lre). Using *PmaIfn1* as a query, we further identified three molecules with more than 90% identity from the full-length transcriptome database of lamprey *Lethenteron camtschaticum*, which were named as *LcIfn1-LcIfn3.* All the identified sequences have been deposited to NCBI database under indicated accession numbers **(Fig. 1B and SI Appendix, Fig. S1A and Dataset S1)**.

**Fig. 1.**
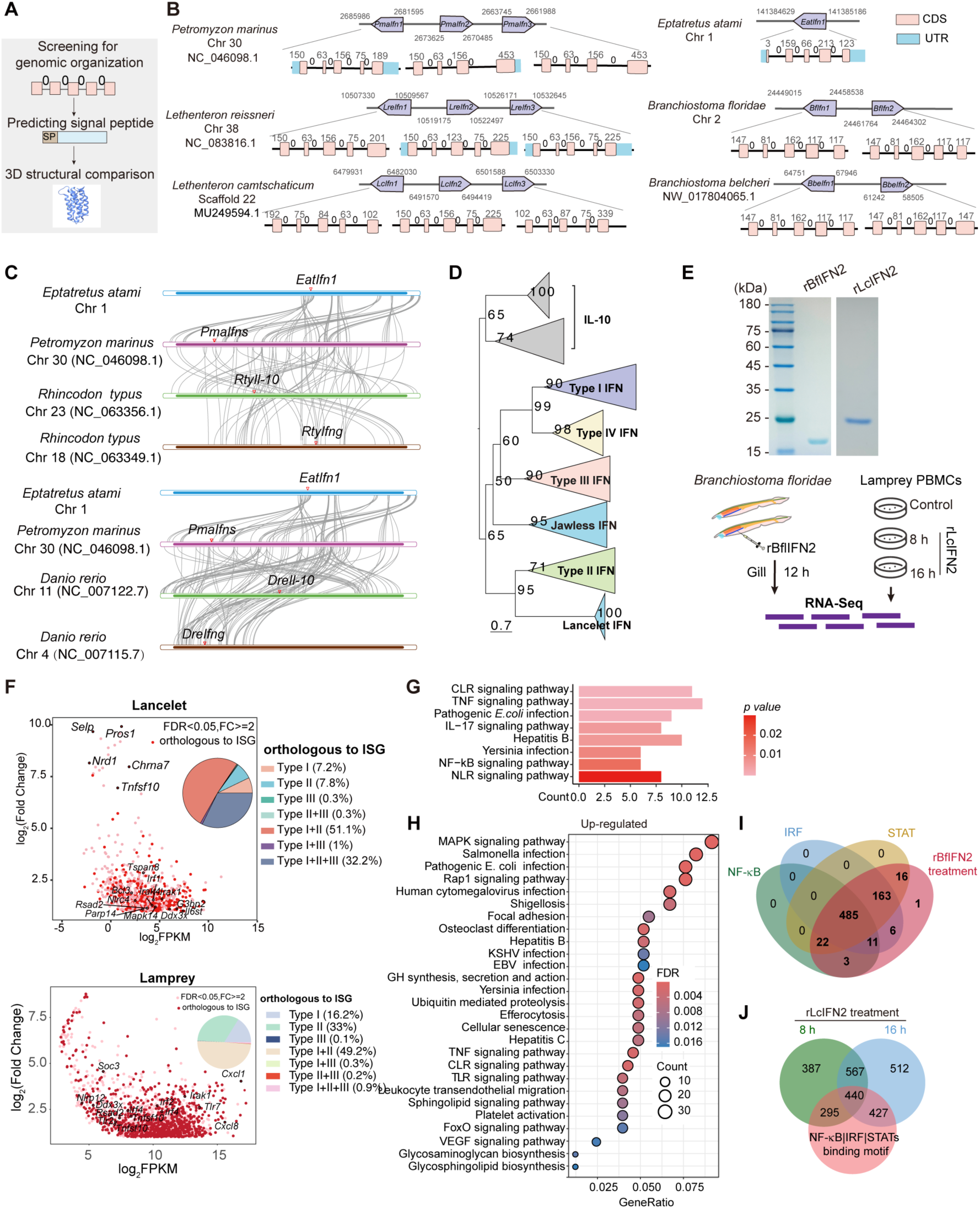
Identification of IFN homologs in agnatha and lancelets. **(A)** Flowchart for identification of the IFN homologs in jawless vertebrates. **(B)** Schematic representation of genomic location, exon-intron organization, and intron phases of the indicated genes from lampreys, hagfish and lancelets. The pink boxes indicated coding regions of exons and lines between boxes indicated introns. The exon length (bp) was numbered upon box, and the intron phase was represented above the line. **(C)** Analysis of the gene linkage and collinearity between hagfish *Eptatretus atami*, lamprey *Petromyzon marinus*, whale shark *Rhincodon typus*, and zebrafish *Danio rerio*. Gray lines indicate genes with collinearity. **(D)** The topology diagram of the phylogenetic analysis of IFNs used IL-10 as outgroup (ML tree). Detailed information for the phylogenetic tree was shown in SI Appendix, Fig. S1D. **(E)** The schematic diagram of stimulation with BfIFN2 and LcIFN2 to adult lancelets and lamprey PBMCs. CBB assay to evaluate the purity of the purified rBfIFN2 and rLcIFN2 from 293T cells. **(F)** Dot plot illustrating the up-regulated genes of lancelet gills and lamprey PBMCs following treatment of rBfIFN2 and rLcIFN2, respectively. The pie chart shows the overlap between the up-regulated genes upon indicated IFNs treatment and those listed in the Interferome database. **(G)** GO analysis of the up-regulated genes in the gills of lancelets following treatment of rBfIFN2. **(H)** KEGG enrichment of the up-regulated genes in response to the treatment of rLcIFN2 protein for 8 h and 16 h. **(I-J)** Venn diagram shows the overlap of the up-regulated genes in response to rBfIFN2 **(I)** or rLcIFN2 **(J)** treatment and the presence of specific transcription factor (TF) binding motifs in their promoter regions.

The identification of putative IFN homologs in jawless vertebrates encouraged us to explore invertebrate genomes **(SI Appendix, Dataset S2)**. Ultimately, two molecules were identified in lancelets *Branchiostoma floridae* (Bf), which were named BfIFN1 and BfIFN2. Additionally, two were identified in lancelets *Branchiostoma belcheri* (Bbe), named BbeIFN1 and BbeIFN2 **(Fig. 1B and SI Appendix, Dataset S1)**. Although the protein sequences of lancelet IFNs exhibit only about 23% identity to those of lamprey IFNs, their predicted structures and genomic organizations strikingly resembled those of vertebrate IFNs **(SI Appendix, Fig. S1A and B)**. Given that the intron numbers and the intron-exon phases of invertebrate IFNs may differ from those in vertebrates, such as *PmaIfn3*, and considering that IFNs are characterized by highly conserved three-dimensional structures despite substantial sequence divergence, we further employed Foldseek program to search for molecules with structurally analogous proteins in invertebrate proteomes using multiple vertebrate IFN structures as queries, and the candidates were further analyzed for signaling peptides **(SI Appendix, Fig. S1C and D)**. Due to the lack of proteins predicted from some evolutionarily critical deuterostome species in AFDB v4 database, we first generated a custom database by extracting protein sequences ≤400 amino acids and predicting their tertiary structures from *Strongylocentrotus purpuratus* (purple sea urchin; GCF_000002235.5, Spur_5.0), *Lytechinus pictus* (green sea urchin; GCF_037042905.1, Lp3.0), and *Styela clava* (tunicate; GCF_013122585.1, ASM1312258v2). Moreover, we incorporated the predicted IFN proteins from lamprey and lancelet into this custom dataset. Subsequently, we constructed a hybrid database by integrating the public AFDB v4 database with our custom-generated predictions. Leveraging this hybrid structural database, we performed a comprehensive screen for proteins structurally homologous to typical IFNs using Foldseek. This analysis successfully identified established IFN homologs in both lampreys and lancelets, yet failed to detect any secreted proteins shorter than 300 aa with significant structural similarity to the canonical IFN fold in other invertebrate lineages **(SI Appendix, Fig. S1C and D)**, thereby supporting the notion that IFNs are restricted to basal chordates and vertebrates.

Additionally, analysis of gene linkage and collinearity not only shows synteny between hagfish and lamprey *Ifn* loci, but also among lamprey *Ifn* homologs and shark *Il-10* and *Ifng*, as well as zebrafish *Il-10* and *Ifng* loci **(Fig. 1C)**, supporting that the identified genes in jawless vertebrates represent ancestral IFN homologs shared with jawed vertebrates. Since IL-10 has been demonstrated to have an evolutionary relation to IFNs, we then constructed a phylogenetic tree using IL-10 as an outgroup. The results revealed that jawless IFN homologs formed a sister group with type I, type III and type IV IFNs, while lancelet IFNs clustered as a sister branch to IFNγ, the ancient IFN subgroup **(Fig. 1D and SI Appendix, Fig. S1E)**.

To functionally validate the newly identified molecules as bona fide IFNs, we selected LcIFN2 from the Japanese lamprey (*Lethenteron camtschaticum*, Lc) as a representative candidate, given that the three LcIFN paralogs share ∼90% intra-group sequence identity and LcIFN2 exhibits the highest similarity to the other two proteins **(SI Appendix, Fig. S1A)**. Likewise, we selected BfIFN2 from lancelet (*Branchiostoma floridae,* Bf) as another representative, considering that BfIFN1 and BfIFN2 also share ∼90% sequence identity **(SI Appendix, Fig. S1A)**. Upon successful cloning, BfIFN2 and LcIFN2 were overexpressed, and the recombinant proteins (rBfIFN2 and rLcIFN2) were purified from 293T cells. The purified proteins were subsequently used for intraperitoneal injection of adult lancelets or for stimulation of lamprey PBMCs **(Fig. 1E)**, followed by RNA-sequencing (RNA-seq) analysis of the treated samples. Differentially expressed genes (DEGs) analysis revealed that the up-regulated genes contained many well-known ISGs like *Rsad2*, *Parp14*, and *Irf1* in both rBfIFN2 treated gill slits and rLcIFN2 stimulated PBMCs **(Fig. 1F)**. Approximately half of the up-regulated genes induced by rBfIFN2 and rLcIFN2 overlapped with those induced by type I and type II IFNs in humans **(Fig. 1F)**. Gene Ontology (GO) enrichment showed that the up-regulated genes upon rBfIFN2 treatment were involved in immune responses activating signaling pathway, pattern recognition receptor signaling pathway, and regulation of inflammatory response **(Fig. 1G)**. Meanwhile, KEGG enrichment and GO analysis showed that the up-regulated genes upon rLcIFN2 treatment were also enriched in human defense responses like against Hepatitis B/C, KSHV, EBV, *Salmonella*, and *Yersinia* infection as well as MAPK (Mitogen-activated protein kinase), TNF, and TLR signaling pathways **(Fig. 1H and SI Appendix, Fig. S1F)**. Furthermore, transcription factor (TF) binding motifs analysis revealed enrichment for IRF, NF-κB, and STAT binding sites in the promoter regions of those up-regulated genes in both lancelet and lamprey **(Fig. 1I and J)**. These results collectively indicated that both BfIFN2 and LcIFN2 were capable of inducing conserved IFN-like responses, thereby validating their functional equivalence to jawed vertebrate IFNs. Thus, these observations traced the origin of IFN back to basal chordates, the transition from invertebrates to vertebrates.

### Lamprey IFNs exert antiviral functions

In jawed vertebrates, IFNs serve as pivotal mediators of antiviral defense by inducing the expression of a broad array of ISGs that act to restrict viral replication (29). Due to the challenges in isolating primary cells from lancelets and the lack of natural viruses of lampreys, to reveal whether IFN-mediated antiviral immunity represents an ancestral and conserved antiviral defense strategy at the transition from invertebrates to vertebrates, we separated Japanese lamprey PBMCs, and then infected them with Vesicular stomatitis virus (VSV)-eGFP, a broad-host-range rhabdovirus widely used as a reporter virus for antiviral assays (30). When compared to PMBCs without infection, VSV-infected lamprey PBMCs gradually become round, shed, and die, typical symptoms of viral infection **(Fig. 2A and B)**. Then, increased VSV-glycoprotein (VSV-G) protein expression was detected in VSV-infected PBMCs by western blot (WB) **(Fig. 2C)**. Among PBMCs, lamprey monocytes were more susceptible to VSV infection than lymphocyte-like cells **(SI Appendix, Fig. S2A)**. Besides PBMCs, we also intraperitoneally injected different PFUs VSV-eGFP into adult lampreys. By 24 hours after injection, a gradual increase in the proportion of monocytes, and a concomitant decrease in the proportion of lymphocytes were observed in PBMCs **(Fig. 2D and SI Appendix, Fig. S2B)**. qPCR assay successfully detected the abundance of viral RNAs in various lamprey tissues upon injection **(Fig. 2E)**, and hematoxylin and eosin (HE) staining revealed the significant pathological changes in the kidneys, accompanied by immune cells infiltration **(Fig. 2F)**. Thus, VSV could successfully infect lampreys, which could be used to investigate the antiviral functions of the newly identified IFNs. After infection with VSV, lamprey PBMCs were subjected to transcriptome sequencing, which identified a set of DEGs consisting of three modules **(Fig. 2G)**. Module I included genes up-regulated upon VSV-eGFP infection at 24 h and 48 h. Module II comprised down-regulated genes upon VSV-eGFP infection at 24 h and 48 h, while Module III contained genes up-regulated at 24 h and down-regulated at 48 h upon VSV infection. All three identified *LcIfns* and a set of ISGs like *Mx1* (Myxovirus resistance 1), *Rsad2* (Radical S-adenosyl methionine domain-containing 2), *Isg20* (Interferon-stimulated exonuclease gene 20), *Isg15* (Interferon-stimulated gene 15), *ly6E* (Lymphocyte antigen 6 complex, locus E) and *Parp12* (Poly(ADP-ribose) polymerase 12) were enriched in Module I **(Fig. 2H and I)**. Notably, after matching the up-regulated genes to the Interferome database, 61.98% was found to be homologous to the known ISGs, such as PRRs like *Rig-I* (Retinoic acid-inducible gene I), *Znfx1* (Zinc finger NFX1-type containing 1) and adaptors like TRAFs (TNF receptor-associated factors) **(Fig. 2I)**. We also used qRT-PCR to confirm the up-regulated abundance of ISGs, such as *Isg20*, *Parp12*, *Rsad2*, and *Mx1* **(SI Appendix, Fig. S2C and D)**. Meanwhile, KEGG enrichment showed that genes in Module I are involved in NLR, TLR, MAPK, CLR and RLR signaling **(Fig. 2J)**, while Module II and Module III consisted of growth hormone synthesis, secretion and action, cGMP (Cyclic guanosine monophosphate, cGMP) -PKG (Protein kinase G, PKG) signaling and Rap1 (Ras-associated protein 1) pathway **(SI Appendix, Fig. S2E and F)**. Thus, these findings indicated that the IFN-based response played a crucial antiviral role in lampreys.

**Fig. 2.**
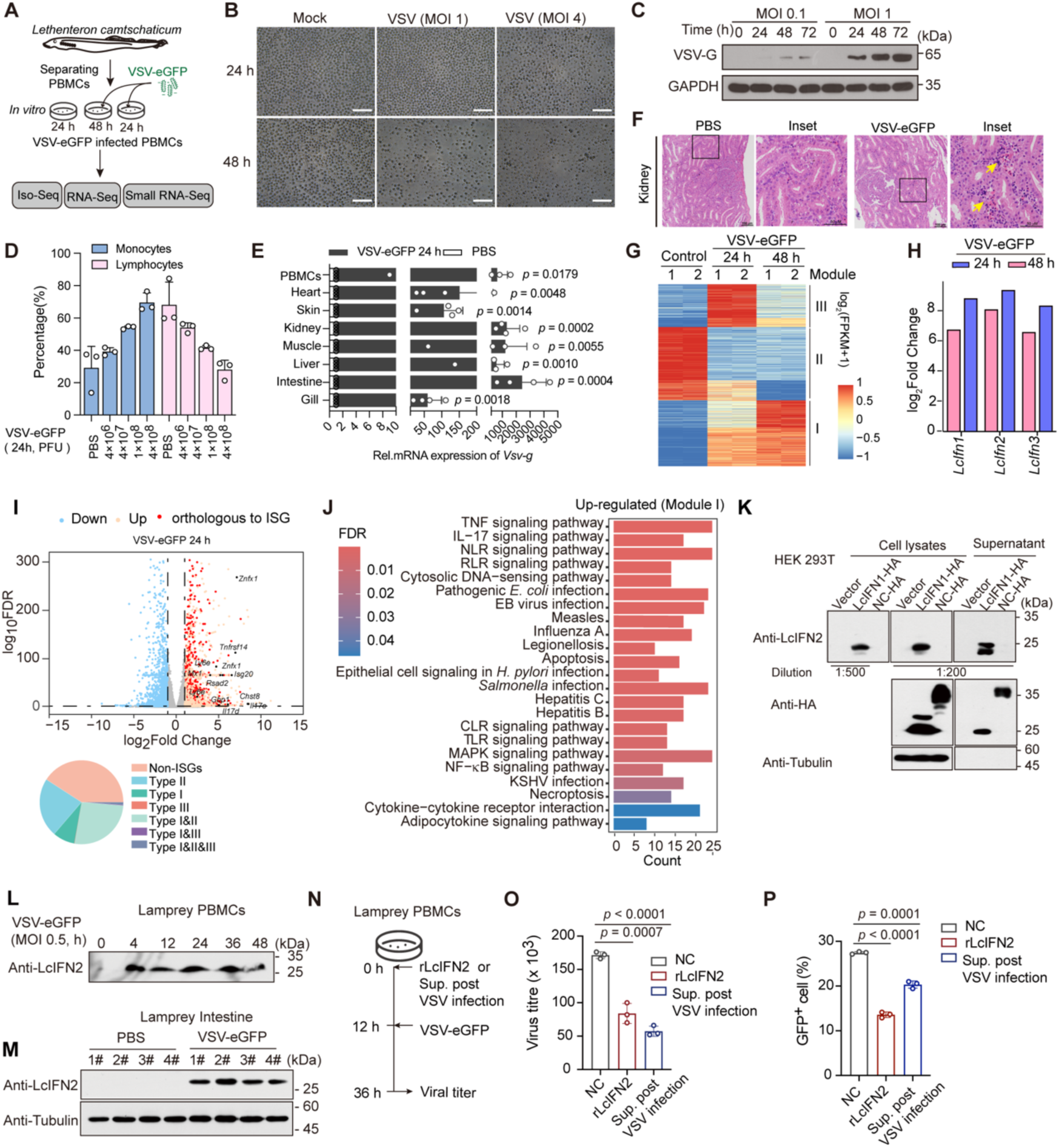
LcIFN2 exerts its antiviral activity. **(A)** The diagram of establishing the VSV-GFP infected lamprey PBMCs. **(B)** The morphology of PBMCs after being infected with VSV-eGFP for 24 h or 48 h. Scale bar, 50 μm. **(C)** Immunoblot of VSV-G in lamprey PBMCs infected with VSV. **(D)** Proportion changes of lamprey PBMC subsets following intraperitoneal injection of VSV-eGFP at different PFUs. All values are means ±SD of three independent experiments. **(E)** Distribution of *Vsv-g* mRNA in various lamprey tissues following intraperitoneal injection of VSV-eGFP (1×10^8^ PFU) determined by qRT-PCR. **(F)** Representative HE staining images of lamprey kidney injection with VSV-eGFP (1×10^8^ PFU) for 24 h. Yellow arrows indicated the presence of immune cell infiltration. **(G)** Heatmap shows the expression profile of lamprey PBMCs infected with VSV-eGFP (MOI = 0.5). **(H)** The mRNA abundance of *LcIfn1*, *LcIfn2* and *LcIfn3* was enhanced upon VSV-eGFP (MOI = 0.5) infection. **(I)** The volcano plot shows the fold changes of genes after VSV-eGFP (MOI = 0.5) infection for 24 h. The pie plot in bottom shows the up-regulated genes of lamprey upon VSV infection for 24 h compared with the Interferome database. **(J)** KEGG pathway enrichment of up-regulated genes after VSV-eGFP infection for 24 h and 48 h. **(K)** WB assays to evaluate the binding specificity and affinity of the anti-LcIFN2 polyclonal serum to the indicated sample. NC-HA is an HA-tagged secreted protein, which serves as negative control. **(L)** WB assays to detect the LcIFN2 protein in culture supernatants of lamprey PBMCs upon VSV-eGFP infection at indicated time points. **(M)** WB assay to detect the LcIFN2 expression in lamprey intestines infected with VSV-eGFP (1×10^8^ PFU) for 24 h. Distinct numbers indicate individual Japanese lampreys under different treatments. **(N)** The schematic diagram of lamprey PBMCs infected with VSV-eGFP. **(O)** The viral titer of VSV-eGFP recovered from lamprey PBMCs with or without treatment of rLcIFN2 or supernatant derived from VSV-infected PBMCs. VSV-eGFP MOI = 0.2. Data represent the mean ±SD of three independent experiments. Student’s *t*-test. **(P)** FACS analysis of GFP positive monocytes in lamprey PBMCs pre-treated with PBS or rLcIFN2 or supernatant derived from VSV infected PBMCs for 16 h followed by VSV-eGFP infection.

To further confirm whether lamprey IFNs could induce antiviral states, we expressed and purified rLcIFN2 from *E. coli* BL21(DE3) **(SI Appendix, Fig. S2G)**, and generated a polyclonal antibody against it. Using the prepared anti-LcIFN2 polyclonal antibody which could recognize LcIFN2 *ex vivo* at the dilution ratio of 1:500 **(Fig. 2K**), we confirmed that LcIFN2 was up-regulated and secreted to cell culture supernatant of lamprey PBMCs upon VSV-eGFP infection **(Fig. 2L)**. Increased abundance of LcIFN2 protein was further observed in lamprey intestines upon intraperitoneal injection of VSV-eGFP **(Fig. 2M)**. Then, both purified rLcIFN2 from 293T cells and filtered supernatant of VSV-infected PBMCs were used to treat lamprey PBMCs for 12 h before VSV-eGFP infection **(Fig. 2N)**. Plaque assay revealed a significant reduction in VSV viral titer upon treatment with rLcIFN2 or viral infection-derived supernatants **(Fig. 2O)**. Consistently, FACS analysis demonstrated a marked reduction in the proportion of VSV-GFP positive cells upon these two treatments **(Fig. 2P)**. Meanwhile, pre-treatment of lamprey PBMCs with rLcIFN2 or with VSV infection-derived supernatants before viral infection resulted in a marked upregulation of the mRNA abundance of *LcMx1*, *LcRsad2* and *Isg20* compared with untreated control **(SI Appendix, Fig. S2H)**. Collectively, these results indicated that LcIFN2 has roles in restricting viral infection, and suggested the common transcriptional regulation of IFNs across species upon antiviral responses.

### CRFB7 and CRFB14 form heterodimeric receptor for lamprey IFN

IFNs used to exert their biological effects by binding to specific receptors (7). IFN receptors belong to the class II cytokine receptor family, also known as cytokine receptor family B (CRFBs). Although identification of class II cytokine receptors has been previously performed by Max D. Cooper and colleagues in sea lamprey (*Petromyzon marinus*) (31), to identify the receptors of LcIFN2, we re-analyzed the composition of class II cytokine receptors in *Lethenteron reissneri* using HMM and BLAST approaches, and then identified seven CRFBs. Given the difficulty in establishing definitive orthologous relationships between these CRFBs and other vertebrate IFN receptors, we named these receptors as CRFB5, CRFB7, CRFB8, CRFB10, CRFB11, CRFB14, and CRFB15 based on the phylogenetic tree, respectively **(SI Appendix, Fig. S3A and B)**. Notably, compared to the previously identified receptors in *Petromyzon marinus*, we newly identified two more CRFBs, CRFB7 and CRFB8 in *Lethenteron reissneri*. As **Fig. S3A** shown, all lamprey CRFBs contain an N-terminal signal peptide, two fibronectin type III (FNIII) domains, including one tissue factor domain and one interfer-bind domain, and a transmembrane (TM) domain, consistent with the structural features of vertebrate IFN receptors **(SI Appendix, Fig. S3A)**. Notably, the two FNIII domains in all lamprey CRFBs are organized in a characteristic of 1-2-1-0-1 intron-phase array, like ancestral class II cytokine receptors (32). Furthermore, gene syntenic analysis suggested that lamprey *Crfb10* exhibits conserved synteny with shark *F3b*, which in turn is syntenic to zebrafish *F3a* and *F3b*, supporting orthology between lamprey *Crfb10/Crfb11* and tissue factor (*F3*). Importantly, genes flanking lamprey *Crfb14* exhibited syntenic conservation with shark *Il10r-Ifnar1* cluster **(SI Appendix, Fig. S3C)**, indicating a close evolutionary relationship between lamprey *Crfb14* and type I IFN receptors.

To identify the receptors of lamprey IFNs, we then performed single-cell RNA sequencing (scRNA-seq) analysis of lamprey *Lethenteron reissneri* PBMCs upon rLcIFN2 treatment, given that Japanese lamprey lacks the chromosome-level assembly of the genome and considering the high protein sequence identity (98%) between LcIFN2 and LreIFN2 **(Fig. 3A)**. It was observed that *Isgs* including *LreMx1*, *LreIsg20*, and *LreParp12* were up-regulated upon rLcIFN2 stimulation for 24 h, indicating that rLcIFN2 can function across distinct lamprey species **(SI Appendix, Fig. S4A)**. Moreover, based on cell markers identified by the previous literature and the analysis of DEGs (33), 15 cell clusters were classified via uniform manifold approximation and projection (UMAP) dimensionality reduction (resolution 0.4) and further categorized into six groups, including the monocytes, neutrophils/DCs, erythrocytes, VLRB^+^ cells, VLRA^+^/C^+^ cells, and MPL proto-oncogene, thrombopoietin receptor (MPL) ^+^ cells **(SI Appendix, Fig. S4B)**. Data analysis showed that the mRNA abundance of *LreMx1*, *LreIsg20* and *LreParp12* was significantly increased in monocytes, neutrophils, VLRB^+^ cells, and VLRA^+^/C^+^ cells **(SI Appendix, Fig. S4A)**.

**Fig. 3.**
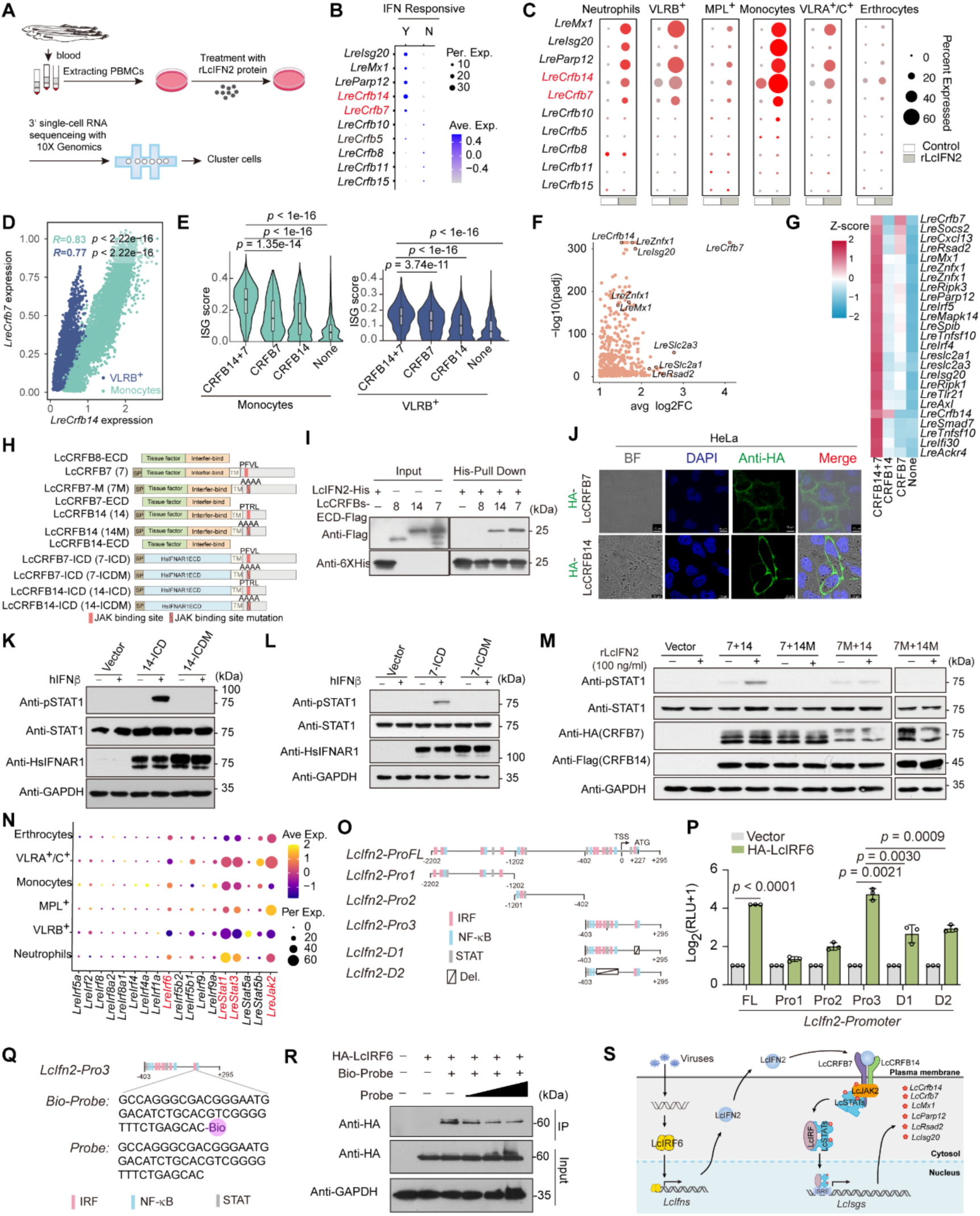
LcCRFB7 and LcCRFB14 form dimeric receptor for LcIFN2. **(A)** Schematic diagram of the experimental design for scRNA-sequencing. **(B)** Dot plot showing the gene expression of the indicated genes across different cell populations following rLcIFN2 treatment. Y indicates an IFN-responsive cluster, while N stands for an IFN-non-responsive cluster. **(C)** Dot plots showing the expression of the indicated genes across various cell types following rLcIFN2 treatment. **(D)** Scatterplots display correlations of *LreCrfb14* and *LreCrfb7* expression in monocytes and VLRB^+^ cells. Blue and green dots indicate VLRB^+^ cells and monocytes, respectively. **(E)** Violin plot showed the ISG score across various cell subpopulations defined by the mRNA abundance of *LreCrfb7* and *LreCrfb14*. CRFB14+7 denotes cells co-expressing *LreCrfb7* and *LreCrfb14*; CRFB14 and CRFB7 denote populations with detectable expression of *LreCrfb14* or *LreCrfb7*, respectively; “None” indicates cells with undetectable expression of both receptors. **(F)** Dotplot showed marker genes in the CRFB7+14 cell subpopulation in monocytes. **(G)** Heatmap illustrating the expression levels of ISGs across distinct monocyte subpopulations. **(H)** Schematic representation of LcCRFBs vector construction. **(I)** His pull-down assay demonstrated that LcIFN2 could interact with LcCRFB14 and LcCRFB7. **(J)** Confocal fluorescence microscopy to detect the localization of LcCRFB7 and LcCRFB14 in HeLa cells. Scale bar, 10 μm. **(K-L)** WB assay to detect the phosphorylated (p-) STAT1 in A549 *^Ifnar1-/-^* cells transfected with indicated vectors, followed by recombinant hIFNβ protein (100 ng/ml) treatment for 30 min. **(M)** WB assay to detect the phosphorylated (p-) STAT1 in HeLa cells transfected with indicated vectors, followed by rLcIFN2 (100 ng/ml) treatment for 30 min. **(N)** Dot plot showed the mRNA abundance of the indicated genes across various cell types. **(O)** Schematic representation of *LcIfn2* promoter regions. **(P)** Luciferase activity analysis to detect the transcriptional regulation of LcIRF6 to the promoter of *LcIfn2* in 293T cells. Data represent the mean ±SD of three independent experiments. Student’s *t*-test. **(Q)** DNA oligonucleotide probes are used for DNA pull-down. The probe was synthesized with a biotin modification at the 5’ end. **(R)** DNA pull-down showed that LcIRF6 could bind to the *LcIfn2* promoter. **(S)** The model of lamprey IFN signaling pathway.

To further determine the receptors of lamprey IFN2, we divided the cells into two clusters based on the fold change of *Isgs* upon rLcIFN2 treatment, one is IFN-responsive, while the other is IFN-non-responsive. We found that *LreCrfb7* and *LreCrfb14* showed high expression in IFN-responsive cluster **(Fig. 3B)**. Among rLcIFN2-responsive cell populations, monocytes and VLRB⁺ cells exhibited the most robust transcriptional responses **(Fig. 3C)**, concomitant with strong co-expression of *LreCrfb7* and *LreCrfb14* within these subsets **(Fig. 3D and SI Appendix, Fig. S4C)**. Then, we subdivided monocytes and VLRB^+^ cells into four distinct subpopulations based on the expression level of *LreCrfb7* and *LreCrfb14*, including cells expressing only *LreCrfb7* (named as CRFB7, counts > 0), only *LreCrfb14* (named as CRFB14, counts > 0), both receptor (named as CRFB14+7, counts > 0), or neither (named as None). Then, we calculated the ISG score by AUCell, which revealed that cells co-expressing *LreCrfb7* and *LreCrfb14* exhibited the highest ISGs induction than cells expressing each receptor (**Fig. 3E-G**). Cells expressing a single receptor exhibited slight ISGs induction, likely attributable to undetectable low-level expression of another receptor below the sensitivity threshold of single-cell sequencing (**Fig. 3E-G**). These analyses suggested that CRFB7 and CRFB14 are likely to be the receptor dimer for lamprey IFN2.

To confirm CRFB7 and CRFB14 are indeed the receptor dimer of lamprey IFNs, we then cloned *LcCrfb8*, *LcCrfb7* and *LcCrfb14* from lamprey *Lethenteron camtschaticum*, which showed over 95% identities with their *Lethenteron reissneri* counterparts **(Fig. 3H)**. After purifying the recombinant proteins from 293F cells and *E. coli*, we used His pull-down assays to confirm that the extracellular domain (ECD) of LcCRFB7 (LcCRFB7-ECD) and LcCRFB14 (LcCRFB14-ECD), but not LcCRFB8-ECD directly interact with LcIFN2 **(Fig. 3I)**. Co-immunoprecipitation (Co-IP) assays further demonstrated that LcCRFB7-ECD can interact with LcCRFB14-ECD (**SI Appendix, Fig. S4D)**. Constantly, structural modeling using AlphaFold3 suggested that LcCRFB7/14 dimer is capable of binding LcIFN2 (**SI Appendix, Fig. S4E)**.

To further reveal whether LcCRFB14 and LcCRFB7 could activate downstream JAK-STAT signaling, we first confirmed their surface expression via flow cytometry and immunofluorescence **(Fig. 3J and SI Appendix, Fig. S4F)**. Sequences analysis further identified conserved JAK binding motifs (PxxL) within the C-terminal intracellular domains (ICDs) of both receptors. Consequently, we constructed a series of expression vectors, including chimeric receptors comprising the extracellular domain (ECD) of human IFNAR1 (HsIFNAR1) fused to the ICDs of LcCRFB7 or LcCRFB14 (named as 7-ICD and 14-ICD, respectively), along with corresponding PxxL motif mutants (named as 7-ICDM and 14-ICDM, respectively) **(Fig. 3H).** After verifying the surface expression of these chimeras by flow cytometry in 293T cells **(SI Appendix, Fig. S4F)**, we then transfected them into A549*^Ifnar1-/-^* cells, following stimulation with HsIFNβ for 30 min. It was observed that constructs of both LcCRFB7-ICD and LcCRFB14-ICD induced the phosphorylation of STAT1, whereas constructs containing mutations of the PxxL motif abolished this ability **(Fig. 3K and L)**, demonstrating that intact JAK-binding motifs are required for signal transduction. To further reveal whether LcCRFB7 and LcCRFB14 could directly bind to LcIFN2 and activate JAK-STAT signaling pathway, HeLa cells were transfected with WT LcCRFBs and their PxxL motif mutants upon stimulation with rLcIFN2 for 30 minutes. As the results demonstrated, rLcIFN2 stimulation activated the phosphorylation of STAT1 when LcCRFB7 was co-transfected with LcCRFB14 **(Fig. 3M)**. Meanwhile, mutations of the JAK-binding motif in LcCRFB7 and LcCRFB14 abolished this ability **(Fig. 3M)**. Thus, we suggested that co-expression of LcCRFB14 and LcCRFB7, and their assembly into a functional heterodimeric complex, is essential for robust IFN induction in lamprey. Notably, both LcCRFB14 and LcCRFB7 were inducible upon rLcIFN2 stimulation **(Fig. 3C)**, suggesting that the initial binding to LcIFN2 to may trigger a weak but basal IFN response, which is subsequently amplified upon co-expression and heterodimeric assembly of these two receptors.

To further reveal the downstream signaling of lamprey IFNs, we performed genomic screening of other key molecules, and found that lamprey possesses thirteen IRFs, one JAK and four STATs (**SI Appendix, Fig. S5A and S5B)**. The phylogenetic analyses of these genes were presented in **SI Appendix, Fig. S5A and S5B**. By analyzing the scRNA-seq data, we found that STAT1, STAT3, IRF6 are widely co-expressed in various cell types in lamprey **(Fig. 3N)**. As **Fig. 3O** demonstrated, the promoter of *LcIfn2* contains several IRF binding motifs **(Fig. 3O)**. Thus, we conducted luciferase reporter assays to detect whether LcIRF6 can drive the transcription of reporter containing *LcIfn2* promoters. We first cloned the promoter sequence of *LcIfn2* and then constructed a series of reporter plasmids containing the WT *LcIfn2* promoter, or the truncated mutants by deleting the IRF binding motifs based on the pGL3 plasmid **(Fig. 3O)**. Then, these reporter plasmids were co-transfected with HA-LcIRF6-expressing plasmids into 293T cells. The results revealed that LcIRF6 enhanced the expression of the respective *LcIfn2-*promoter containing reporters. However, deletion of the core IRF binding sites in the *LcIfn2* promoter inhibited the expression of reporter gene **(Fig. 3P)**. Furthermore, a biotin-labeled DNA probe was constructed and DNA pull-down assays were performed to reveal that LcIRF6 can directly bind to *LcIfn2* promoter **(Fig. 3Q and R)**. Therefore, upon viral infection, lamprey IRF6 can drive the transcription of *Ifn2*, which then binds to its heterodimeric receptor complex containing CRFB7 and CRFB14 to recruit the JAK-STAT signaling to activate the transcription of a series of *Isgs* in lamprey, one of the representatives of jawless vertebrates **(Fig. 3S)**.

### Lamprey IFN orchestrates both innate and adaptive immune responses by reshaping the cytokine and chemokine networks

As key antiviral cytokines induced by viral infection, IFN is a pleiotropic cytokine family that plays pivotal roles in the immune response (2, 6, 34). Unlike adaptive immunity that mediated by T/B cells in jawed vertebrates, jawless vertebrates have an alternative adaptive immune system based on their distinct VLRA-F receptors (35, 36). As shown in **Fig. 3C**, lamprey CRFB7 and CRFB14 were co-expressed in distinct lamprey PBMCs, especially monocytes and VLRB^+^ cells. The significant increase in the proportion of cluster 1 VLRB^+^ cells following high expression of ISGs upon rLcIFN2 stimulation implied the functional involvement of IFN signaling in coordinating both innate and adaptive-like immune responses in jawless vertebrates **(SI Appendix, Fig. S6A)**.

To further elucidate the multiple regulatory functions of LcIFN2 beyond antiviral defense, we first compared the data obtained from rLcIFN2 stimulated with VSV-infected PBMCs. Our results revealed that approximately 25% of the genes exhibited similar expression patterns upon IFN induction or viral infection **(Fig. 4A)**. KEGG enrichment analysis indicated that genes induced by rLcIFN2 stimulation at 8 h and 16 h, as well as by VSV infection at 24 h and 48 h, are involved in key signaling pathways related to viral infection, including TNF, NLR, RLR, and TLR pathways, among others **(Fig. 4A)**. In contrast, genes that were only up-regulated at 8 h by rLcIFN2 stimulation were associated with pathways related to cell development and differentiation, such as MAPK, Rap, and Hippo signaling pathways **(Fig. 4B)**. Consistently, GSEA analysis of DEGs in monocytes and VLRB⁺ cells following rLcIFN2 stimulation demonstrated significant enrichment not only in the interferon signaling pathway, but also in multiple cytokine-driven pathways such as IL-2 and IL-6 signaling pathways **(SI Appendix, Fig. S6B and C)**. Genes that were commonly up-regulated upon rLcIFN2 stimulation in all cell populations were enriched in defense response to virus and regulation of innate immune response, whereas genes specifically upregulated in monocytes were significantly enriched in pathways related to myeloid cell homeostasis, fatty acid oxidation, and lipid oxidation **(Fig. 4C)**. Meanwhile, genes specifically upregulated in VLRB^+^ cells were enriched in store-operated calcium entry, T cell differentiation **(Fig. 4D)**. These observations indicated that LcIFN2 may play broader immunomodulatory roles beyond antiviral defense.

**Fig. 4.**
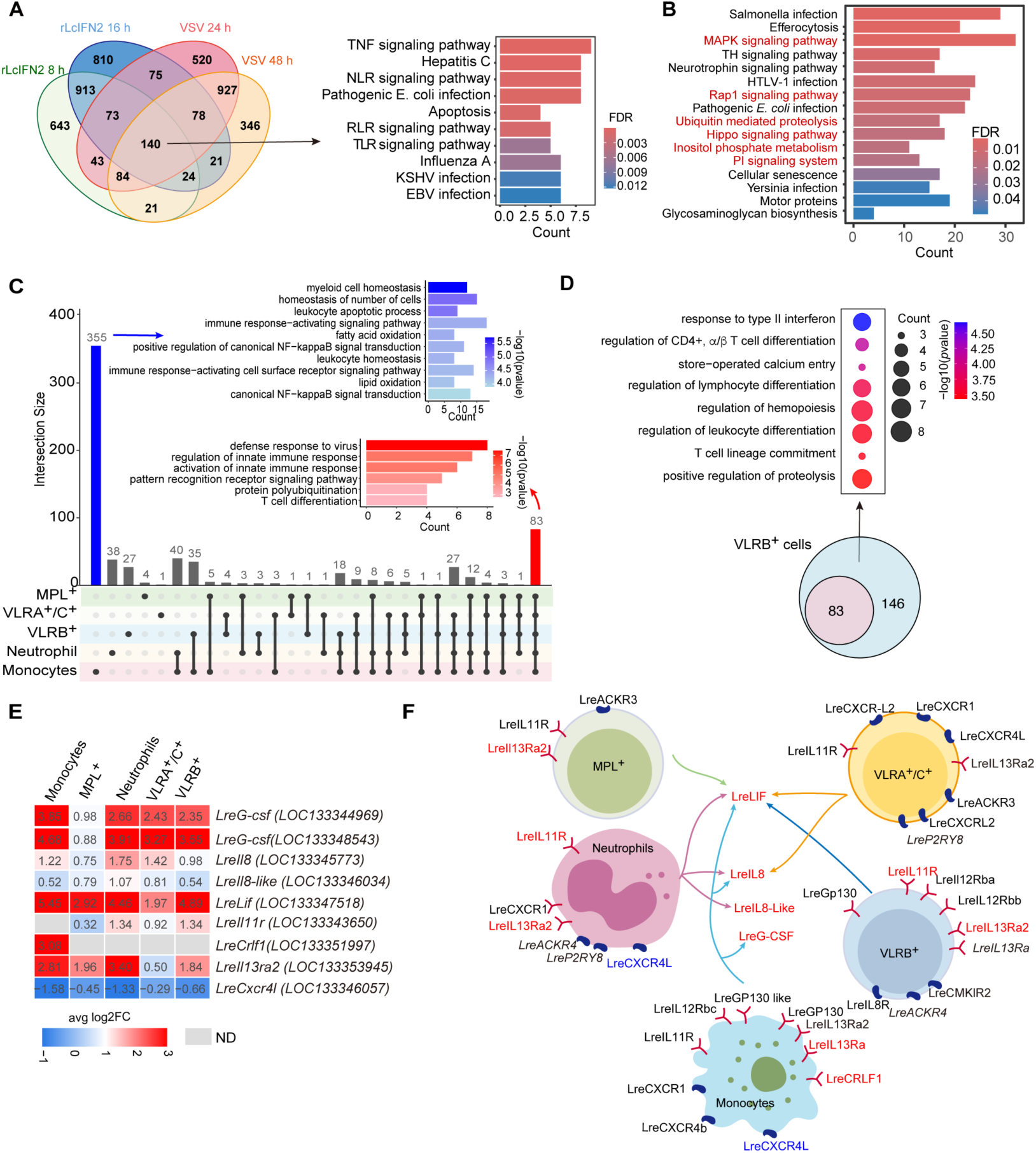
LcIFN2 promotes cellular communications by reshaping the cytokine and chemokine networks. **(A)** Venn diagram illustrating the overlap of the up-regulated genes upon VSV-eGFP (MOI = 0.5) infection for 24 h or 48 h, and rLcIFN2 (100ng/ml) stimulation at 8 h and 16 h. KEGG enrichment analysis of genes up-regulated by both viral infection and rLcIFN2 stimulation. **(B)** KEGG enrichment of genes up-regulated in 8 h upon rLcIFN2 treatment but not VSV infection. Terms highlighted in red indicate pathways associated with cell development and differentiation. **(C)** Upset plot showed the up-regulated genes in response to rLcIFN2 treatment across different cell types. Different colors represent distinct cell types, with gray dots indicating significantly up-regulated genes in the corresponding cell populations. Connecting lines highlight genes that are co-up-regulated across multiple cell groups. The numbers on the bars indicate the count of up-regulated genes. Blue denotes genes exclusively up-regulated in monocytes along with their GO analysis, while red denotes genes commonly up-regulated across all five cell clusters, accompanied by their GO enrichment results. **(D)** KEGG pathway enrichment analysis of genes specifically upregulated in VLRB^+^ cells. Number 83 indicated genes that were commonly upregulated across multiple cell types, highlighted in red panel in Fig. 5C. **(E)** Heatmap showed the fold change of indicated genes upon rLcIFN2 treatment. Detailed expression profiles of more cytokines and putative receptors upon rLcIFN2 induction were presented in SI Appendix, S6D. **(F)** Schematic representation of cell-cell communication mediated by cytokines, chemokines, and their receptors among lamprey PBMCs under rLcIFN2 treatment.

In jawed vertebrates, IFNs are endowed with diverse regulatory effects of immune response by reshaping the cytokines and chemokines network to coordinate both innate and adaptive immunity (37). To reveal how LcIFN2 plays its multifaced roles, we then re-analyzed the DEGs in each cell type upon rLcIFN2 treatment with a focus on the mRNA abundance of cytokines and chemokines and their respective receptors (**Fig. 4E and SI Appendix, Fig. S6D**). As depicted in **Fig. 4F**, rLcIFN2 stimulation induced the upregulation of several cytokines in monocytes and neutrophils, including *G-csf* (Granulocyte colony-stimulating factor), *Il-8* (Interleukin-8), and *Lif* (Leukemia inhibitory factor). In VLRA^+^/C^+^ cells, the expression of *Il-8*, and *Lif* were also increased. Notably, chemokine receptors including *Ackr4* (Atypical chemokine receptor 4), *Cmklr2* (Chemerin chemokine-like receptor 2), and *Cxcl4L* (C-X-C motif chemokine ligand 4-like 1) were expressed in VLRB^+^ cells, whereas *Cxcr-l2* (C-X-C motif chemokine receptor l2) was specifically detected in VLRA^+^/C^+^ cells (**Fig. 4F and SI Appendix, Fig. S6D**). These dynamic expression changes suggested that lamprey IFN may reshape the cytokine and chemokine networks of lamprey, to orchestrate the intercellular communication among distinct cell populations. For instance, the up-regulation of IL-8 in monocytes and neutrophils, coupled with the expression of its receptors IL-8R on VLRB^+^ cells, suggests that LcIFN2 may modulate VLRB^+^-mediated immunity via monocytes and neutrophils-derived IL-8 **(Fig. 4E-F and SI Appendix, Fig. S6D)**. Similarly, the expression of IL-11 in monocytes, along with the increased expression of their receptors IL-11R on VLRB^+^ cells **(Fig. 4E-F and SI Appendix, Fig. S6D)**. Thus, the IFN response in lamprey not only directly activates both innate and adaptive immune cells to combat infections, but also orchestrates their cellular communication and metabolic activities to support sustained immune functions.

### Substitutions in residues essential for Dicer mediated dsRNA processing were coincided with the emergence of the IFN system

Since RNAi and IFN-mediated antiviral mechanisms are two key defense strategies that hosts have evolved, to further understand how IFN based antiviral mechanism gradually became a predominant antiviral response during the evolution of vertebrates, we performed small RNA sequencing of lamprey PBMCs upon VSV-infection. Analysis revealed that while VSV-derived small RNAs (vsiRNAs) were generated from both negative and positive strands and predominantly measured 22 nt **(Fig. 5A and B)**, they accounted for only 0.005% of total small RNAs following VSV infection in lamprey **(SI Appendix, Fig. S7A)**. Such a phenomenon has been previously reported in zebrafish and differentiated mammalian cells like 293T (9, 11) This functional comparison indicated that the antiviral RNAi activity is limited in the entire vertebrate lineage. To reveal the functional alteration of RNAi antiviral mechanism during evolution, we then conducted functional comparison of lancelet and lamprey Dicer proteins. First, sequences screening confirmed that both lancelet and lamprey harbor key components which are essential for miRNA biogenesis, such as Dicer1 and TRBP2, while lack Dicer2 and R2D2, two key proteins involved in siRNA biogenesis in fruit flies **(Fig. 5C-E and SI Appendix, Fig. S7B)**. Given that Dicer-mediated cleavage of long dsRNA depends on RNA binding, ATP hydrolysis-driven translocation along dsRNA, and subsequent processing into ∼22-nt siRNAs (38–40), *in vitro* dsRNA cleavage assays and translocation assays were then conducted to compare the function of lancelet and lamprey Dicer. The results showed that BfDicer but not LcDicer could efficiently cleave dsRNA **(Fig. 5F)**. Consistently, streptavidin displacement assays demonstrated that BfDicer rather than LcDicer was able to translocate along dsRNA **(Fig. 5G and H)**, suggesting that Dicer is functionally changed when invertebrates evolved to vertebrates. Sequence alignment further reveals that two residues V109 and G132 in BfDicer have changed as E/A and S/N in vertebrate Dicer **(Fig. 5I and SI Appendix, Dataset S3)**. Since residues V109 and G132 have been demonstrated to be crucial for dsRNA binding in DmDicer 2 (38), we then mutated these two residues in BfDicer as the corresponding residues in lamprey and human Dicer, which were named as M1(V109E, G132S) and M2(V109A, G132N), respectively. M3 (K76A, D1374A, D1679A) was constructed as a negative control **(Fig. 5J)**, as these residues have been demonstrated to be important for the enzymatic activity of Dicer (39, 41). *In vitro* cleavage assays revealed that BfDicer-M1, M2, and M3 all failed to efficiently cleave dsRNA **(Fig. 5K)**. Supporting this, streptavidin displacement assays revealed that BfDicer-M1, M2 and M3 all disrupted dsRNA translocation **(Fig. 5L)**. Thus, during vertebrate evolution, amino acid substitutions in Dicer that are responsible for dsRNA binding impaired its dsRNA translocation capability, thereby abolishing its ability to cleave dsRNA and execute RNAi-mediated antiviral defense. Additionally, rLcIFN2 stimulation significantly upregulated multiple nucleases and RNAi-suppressing factors, including members of the poly (ADP-ribose) polymerase (PARP) family and ISG20 **(Fig. 5M)**. Given that functional impairment of RNAi has also been attributed to competition for cytoplasmic dsRNA with RIG-I-like receptors (RLRs) (42), these observations suggested that functional antagonism between RNAi and IFN-mediated antiviral pathways represents a widespread and evolutionarily conserved feature across vertebrate species.

**Fig. 5.**
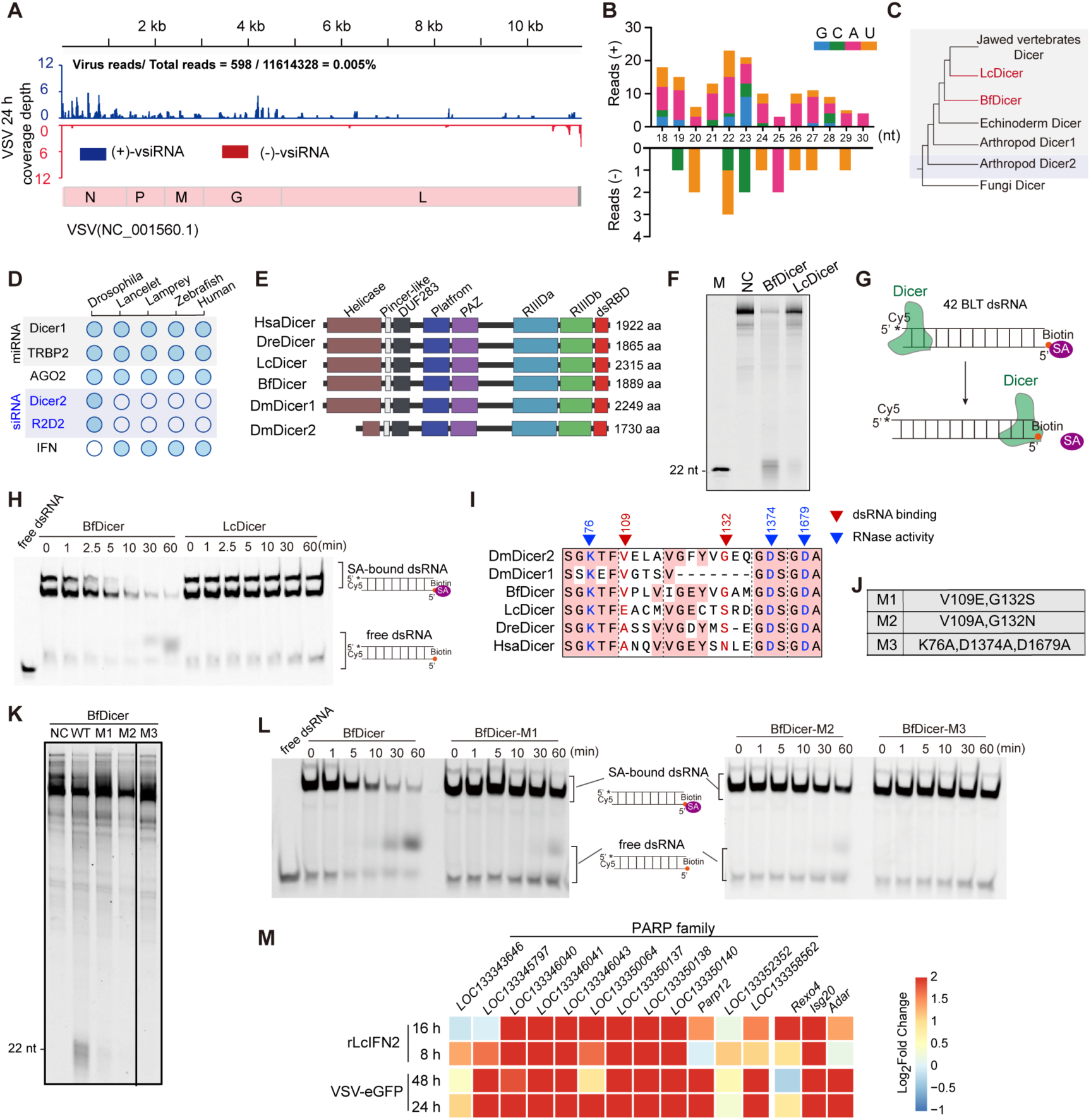
Comparative analysis of components involving in RNAi antiviral mechanism across species. **(A-B)** Distributions of 21- and 23-nt reads along VSV (+) and (-) strands after small RNA deep-sequencing of RNA from lamprey PBMCs infected by VSV-eGFP (MOI = 0.5) for 24 h. **(C)** Summary phylogenetic tree of Dicer homologs. **(D)** Composition of the components involved in RNAi antiviral mechanism across species. **(E)** Schematic diagram of domain architecture of Dicer proteins across species. **(F)** *In vitro* dsRNA cleavage assay showing that BfDicer was capable of cleaving dsRNA. **(G)** Schematic illustration of the streptavidin displacement assay. 42-bp blunt-ended dsRNA (42 BLT dsRNA) was used as substrate, with a biotin moiety (orange circle) covalently linked to its 3’ terminus. Streptavidin (SA, purple oval) binds specifically to the biotin. The Dicer protein (green) initiates binding to dsRNA and then translocates along the duplex toward the 3’ end. Upon reaching the terminus, Dicer physically displaces the streptavidin from the biotin. **(H)** The streptavidin displacement assay showed that BfDicer was capable of translocating along dsRNA. **(I)** Sequence alignment of Dicer proteins across species showing the residues that may affect the activity of Dicer. Red highlights indicate residues involved in dsRNA binding, while blue highlights indicate residues within the catalytic center. Regions with sequence identity higher than 50% are shaded in pink. **(J)** The constructed mutants of BfDicer. **(K)** *In vitro* dsRNA cleavage assay demonstrated that replacements of V109 and G132 in BfDicer as the corresponded residues in lamprey and human Dicer abolished its dsRNA cleavage activity. **(L)** The streptavidin displacement assay showed that the M1, M2, and M3 mutants of BfDicer were unable to translocate along dsRNA. **(M)** Heatmap showed changes in mRNA abundance of indicated genes in response to VSV-eGFP infection or rLcIFN2 stimulation in lamprey PBMCs, presented as log2 Fold change.

## Discussion

### Origin and the evolutionary trajectory of vertebrate IFNs

IFNs constitute an ancient cytokine family that plays an essential role in establishing antiviral states by inducing a set of ISGs, and bridging innate and adaptive immunity via dendritic cell maturation, macrophage activation, and enhanced antigen presentation (6, 34). Although IFNs were previously dated to over 450 million years ago in early jawed vertebrates (17, 21, 22), the present study extends their evolutionary timeline to basal chordates through the identification of homologs in lancelets and jawless vertebrates. Additionally, our functional comparison of Dicer between amphioxus and lamprey supports an evolutionary trade-off wherein RNA-dependent antiviral immunity attenuated concomitantly with the emergence of IFN-mediated defense, indicative of a system-level transition in antiviral defense strategies during vertebrate origin.

Beyond the long-standing mystery surrounding their evolutionary emergence, IFNs display lineage-specific diversification patterns throughout vertebrate phylogeny, especially IFN-Is which exhibit remarkable gene expansion and dynamic copy number variation (CNV) across species (43). Another important phenomenon is that IFN-Is can be classified into two subgroups due to their genomic organization, one is intronless and the other is intron-containing (17). The first intronless IFN likely originated through retrotransposition in amphibian (17), while unequal crossing-over subsequently contributed to the expansion and CNV of multiple type-I IFN copies in specific mammals (43, 44). Notably, CNV also occurs in intron-containing IFNs, particularly evident in teleost fish, such as *Salmo salar*, *Oncorhynchus mykiss*, and *Danio rerio*, where type I IFNs display lineage- or species-specific expansions **(SI Appendix, Fig. S7C and Dataset S4)**. Moreover, the expanded *Ifn* copies in teleost fish often display opposite transcriptional directions and across chromosomes, implying that additional mechanisms beyond unequal crossing-over may also contribute to the IFN diversification in the evolution of early vertebrates. Given that DNA transposons usually lead to regional genomic instability and drive the frequent insertion of duplicated genes, we re-surveyed the *Ifn* loci in several presentative bony fish and identified numerous TIR sequences in their *Ifn* loci. Then, we performed in-depth analysis of rainbow trout *Oncorhynchus mykiss* genome, which harbors an extraordinarily complex type I IFN system. After revealing *Oncorhynchus mykiss* type I IFN genes are distributed across Chr 12, 13, and 16, we then identified several hAT and TcMar TIR in the vicinity of type I IFNs, along with Gypsy and ERV LTRs, as well as LTR/Copia in *Ifng* **(SI Appendix, Fig. S7D and Dataset S4)**. Upon finding numerous TIR sequences surrounding the *Ifn* loci, complete *Tc1* transposons in the rainbow trout *OmyIfn* regions on chromosomes 13 and 16 were identified, including a pair of 5’ and 3’ TIRs, a Tc1 transposase coding sequence, and the target site duplications (TSD) **(SI Appendix, Fig. S7D)**. Given that Tc1 is a member of the *Tc1/mariner* (*TcMar*) superfamily which exhibits broad taxonomic distributions across vertebrates, the identification of contact *Tc1* transposons in *OmyIfn* regions implied the contribution of transposable elements to lineage-specific expansion and diversification of intron-containing IFNs throughout evolution of early vertebrate, in addition to previously demonstrated whole-genome duplication (WGD), tandem duplication, retrotransposition, and non-allelic homologous recombination (NAHR) (17, 43–45).

### The sophisticated function of the ancestral IFNs in immune regulation

Beyond tracing the evolutionary origin of IFN to basal chordates, the present study further revealed that antiviral activity by inducing a set of ISG expression constitutes the primary ancestral function of these ancient signaling proteins. Due to the arm race between viruses and hosts, IFNs in jawed vertebrates have undergone extensive gene duplication and functional diversification, giving rise to four types of IFNs and their corresponding receptors. In mammals, IFN I-IV play specialized roles in host defense, immune regulation, and tissue homeostasis in jawed vertebrates (2). For example, type I, III, and IV IFNs primarily mediate antiviral immunity, while type II IFNs, such as IFN-γ produced mainly by NK cells and activated T cells, function as a central immunoregulatory cytokine and are essential for defense against intracellular bacteria and parasites (7, 46). Specifically, upon initiating acute, cell-intrinsic antiviral response, IFN can precisely coordinate both innate and adaptive immune responses by reshaping cytokine and chemokine networks (3, 46). For examples, upon HSV-1 infection, IFN-α/β can trigger tissue-resident macrophages and DCs to produce CCL2, thereby recruiting inflammatory monocytes (47). IFN-α/β also enhances the expression of MHC molecules and co-stimulatory molecules such as CD80 and CD86 in committed DCs, thereby boosting their ability to stimulate T cells (48). Additionally, human IFN-α/β can promote the differentiation of T cells into IFNγ-producing Th1 cells (49) and activate the antibody responses in B cells, including class switch recombination and IgG subtype switching (50). As the ancestral IFN, here we found that lamprey IFNs not only exhibit conserved antiviral functionality by inducing a set of antiviral ISGs, but also play key immunomodulatory functions by reshaping the cytokines/chemokines networks to coordinate both innate immune cells and VLR cells, bridging the gap between the VLR-based adaptive immune system of jawless vertebrates and the immunoglobulin (Ig)-based system of jawed vertebrates, despite their divergence. Thus, further exploration of the roles of these ancient IFNs in immune cell development or functional regulation may provide valuable insights into the evolution of vertebrate immune systems, especially for the emergence of adaptive immunity.

## Materials and Methods

### Animals

Adult lamprey specimens of *Lethenteron camtschaticum* and *Lethenteron reissneri* were collected from the Wusuli River in Heilongjiang Province, China, and subsequently maintained in laboratory conditions. Lancelets *Branchiostoma floridae* were kindly provided by Prof. Guang Li from Xiamen University (Xiamen, China). All relevant procedures of animal experiments were conducted according to the guidelines established by the Ethics Committee of Sun Yat-Sen University.

### Cells, reagents, and viruses

HEK293T, A549, and HeLa cell lines were purchased from the American Type Culture Collection (ATCC) and maintained in our laboratory. All cells were cultured in Dulbecco’s Modified Eagle Medium (DMEM, Gibco), supplemented with 10% fetal bovine serum (FBS, Gibco), at 37°C with 5% CO₂. 293F cells were purchased from Thermo Fisher Scientific and maintained in our laboratory. 293F cells were cultured in 293F Hi-exp (Shanghai OPM Biosciences) at 37°C with 8% CO₂. The A549 *IFNAR1* knockout cell line was constructed using CRISPR-Cas9-mediated genome editing using gRNA TCATTTACACCATTTCGCAA. CRISPR plasmids were transfected into A549 cells using JetPRIME (Polyplus). GFP-positive single-cell colonies were isolated and subsequently validated at the protein level by Western blot analysis. VSV-eGFP was preserved in our laboratory. VSV-eGFP was propagated and its titers were determined on Vero cells. Antibodies used in this study were listed in SI Appendix, Dataset S5.

### Isolation of lamprey PBMCs

Lamprey PBMCs were isolated using mouse peripheral blood lymphocyte isolation kit (Solarbio), according to the manufacturer’s instructions. Briefly, adult lampreys were anesthetized in MS222 (0.1 g/l; Sigma-aldrich), sterilized with 75% alcohol. Whole blood was collected by tail amputation into EDTA-K2 anticoagulant tubes. The collected blood was diluted to a total volume of 3 ml using the provided blood and tissue diluent. This diluted blood was carefully layered over 3 ml of a separation solution. After density gradient centrifugation, the PBMCs layer was harvested and washed three times with washing buffer. The isolated PBMCs were then cultured in L-15 medium (Gibco) supplemented with 20% FBS at 18°C.

### IFN stimulation and virus infection

To establish the VSV infection model, lamprey PBMCs were separated, cultured in L15 medium containing 20% FBS, and then infected with VSV-eGFP at the indicated MOI. For *in vivo* infection, adult lampreys were anesthetized with MS-222, injected intraperitoneally with VSV-eGFP at the indicated PFUs, and then cultured for 24 h. For preparation of VSV-infected supernatants, lamprey PBMCs were infected with VSV-eGFP (MOI = 0.5) for 12 h. Cell culture supernatants were harvested and centrifuged at 1000 × g to remove cells. The supernatants were subsequently passed through a 0.45 μm filter to eliminate residual cell debris. The filtrates were then subjected to ultrafiltration using 100 kDa cut-off centrifugal filter units. The flow-through fraction was collected and further sterilized by filtration through a 0.22 μm filter before use.

For *in vitro* stimulation, the isolated lamprey PBMCs were maintained in L15 medium containing 20% FBS and treated with recombinant LcIFN2 (rLcIFN2) for 8 or 16 h, while the control group received a corresponding volume of sterile PBS.

For rBfIFN2 treatment in adult lancelets, *Branchiostoma floridae* were anesthetized with MS-222 (0.1 g/l) and injected intracoelomically with 200 ng of rBfIFN2 in 30 μl PBS. Control animals received an equal volume (30 μl) of sterile PBS. After injection, animals were maintained in 0.22 μm-filtered seawater at room temperature for 12 h. Following incubation, lancelet individuals were dissected under a stereomicroscope, and gills were collected for RNA extraction. Each experimental group consisted of two distinct individuals.

### Full-length transcriptome sequencing and assembly (PacBio Iso-Seq)

For full-length transcriptome analysis, high-quality total RNA was extracted as above described. Libraries for full-length transcriptome sequencing were prepared using the SMRTbell Express Template Prep Kit 2.0 (Pacific Biosciences) according to the manufacturer’s instructions. Raw subreads were processed using the IsoSeq3 pipeline (SMRT Link software) to generate high-quality full-length transcripts. Redundant isoforms were removed by CD-HIT, and the non-redundant transcriptome was used for downstream analysis. Functions of transcripts were annotated by alignment to public databases, including NR, SwissProt, GO, and KEGG, using BLAST and Diamond.

### Structural simulation, phylogenetic analysis, and transcription factor binding motif prediction

The 3D structures of proteins were predicted by ESMFold (51). Additionally, the LcIFN2 and LcCRFB7/14-ECD complex was modeled on the AlphaFold3 online server (https://alphafoldserver.com/) (52). The predicted structures were then aligned and compared using Foldseek (V9.4.27) with the easy-search module (53). The parameters were set as follows: E-value cutoff = 1, TM-score ≥ 0.4, query coverage ≥ 0.7, and target coverage ≥ 0.5. Domain structures were analyzed by performing the Pfam program (http://smart.embl-heidelberg.de/). Multiple sequence alignment of IFN homologs was performed using ClusterW, and the maximum likelihood phylogenetic tree was constructed by IQ-TREE (54) with the best-fit substitution model determined by the BIC criterion. Node support was evaluated using 5,000 bootstrap replicates. The phylogenetic trees were visualized and managed using the evolview tool (https://www.evolgenius.info/evolview/). The gene promoter sequences (2 kb upstream of the coding sequence) were extracted using TBtools (55). Promoter motifs scanning were performed using FIMO (56), with known transcription factor binding motifs obtained from the JASPAR database (57). Matches with a *P*-value < 1e-5 were considered significant.

### Single cell RNA-seq library preparation and data analysis

The single cell RNA-seq libraries were prepared using the Chromium Single Cell 3’ Library & Gel Bead Kit v3 (10 × Genomics) according to the manufacturer’s protocol. Briefly, cell suspensions were processed for droplet generation, followed by emulsion breakage, bead collection, reverse transcription, and cDNA amplification to generate the final libraries. Sequencing was performed on an Illumina platform at BeiRui (Beijing, China).

Raw sequencing reads were processed using Cellranger 9.0 and aligned RNA to the *L. reissneri* genome. Then the generated gene expression matrix was processed and analyzed using the Seurat package (v5.1.0) in R (V4.3). Cells with fewer than 200 expressed genes or with >5% mitochondrial gene content were filtered out. Genes detected in fewer than 3 cells were also excluded. The data were normalized using the NormalizeData function with the default log-normalization method. Next, different datasets were integrated with the harmony algorithm following the RunPCA and RunHarmony commands with default parameters. Highly variable genes were identified using FindVariableFeatures, and data were scaled using ScaleData. Cell clustering was conducted using the FindNeighbors and FindClusters functions with a resolution of 0.4. UMAP was performed using RunUMAP for visualization of the cell clusters. Marker genes for each cluster were identified using the FindAllMarkers function based on the *Wilcoxon rank-sum* test. Cell types were annotated by comparing cluster-specific marker genes with known canonical markers from published literature and reference databases. Differential gene expression analysis was conducted using the FindMarkers function.

### Generation of anti-LcIFN2 polyclonal antibody

The coding sequence of LcIFN2 without the signal peptide was cloned into the pET-28a vector and expressed in *Escherichia coli* BL21 (DE3) cells. Protein expression was induced by 1 mM IPTG at 37 °C for 16 h. Cells were harvested and lysed by sonication. Since the recombinant protein was expressed in inclusion bodies, it was purified using inclusion body isolation. The purity of the recombinant LcIFN2 protein was confirmed by SDS-PAGE analysis. Polyclonal antibody against LcIFN2 was custom-produced by GenScript (Nanjing, China) using purified rLcIFN2 as the immunogen. Briefly, New Zealand white rabbits were immunized with purified rLcIFN2 emulsified in Freund’s complete adjuvant. Serum was collected following immunization and affinity-purified using antigen-specific affinity chromatography. The specificity of the purified antibody was validated by western blot against rLcIFN2.

### dsRNA preparation

Long dsRNA was prepared using the first 200 nt of the GFP coding sequence. Sense and anti-sense strands were synthesized by *in vitro* transcription with T7 RNA polymerase (T7 MEGAscript kit, Ambion). RNA was labelled by incorporation of 1/10th of Cy3-UTP (Beyotime) in the IVT reaction. Following Turbo DNaseI treatment (Invitrogen), the single-stranded RNA (ssRNA) was purified using Beckman magnetic beads. The purified ssRNA was annealed in annealing buffer (10 mM Tris-HCl, pH 8.0, 20 mM NaCl) using the following program: 95°C for 3 min, cooled to 75°C and incubated for 30 min, then cooled at a rate of -1°C/min to 4°C. The ssRNA was removed by RNaseI treatment (Invitrogen) and purified using TRIzol reagent according to the manufacturer’s instructions.

### Purification of recombinant Dicers and *in vitro* dicing assay

Recombinants BfDicer and LcDicer were purified from 293F cells. Cells were transfected with PLVX-IRES-Puro encoding Flag-tagged proteins and lysed in lysis buffer (50 mM Tris-HCl (pH 7.5), 150 mM NaCl, 3 mM MgCl₂, 5% glycerol, 0.5% Triton X-100, protease inhibitors, Roche) and sonicated. The lysate was centrifuged at 12,000 g for 10 min, filtered through a 0.45 μm membrane, and incubated with FLAG-M2 affinity resin at 4°C overnight. The resin was washed three times with lysis buffer, twice with wash buffer 1 (50 mM Tris-HCl (pH 7.5), 150 mM NaCl, 3 mM MgCl₂, 5% glycerol, 0.1% Triton X-100, 1 mM PMSF), and twice with wash buffer 2 (50 mM Tris-HCl (pH 7.5), 150 mM NaCl, 3 mM MgCl₂, 5% glycerol, 1 mM PMSF). Proteins were eluted by incubating the resin with wash buffer 2 containing 3×FLAG peptide (300 ng/μL) for 40 min. The eluted protein was collected, aliquoted, and stored at -80°C. *In vitro* dicing assay was performed in a 50 μL volume containing recombinant 200 nM Dicer proteins and 50 nM dsRNA in dicing buffer (30 mM Tris-HCl (pH 6.8), 50 mM NaCl, 3 mM MgCl₂, 5% glycerol, 1 mM DTT, 2 U/μL RNasin, Promega). Reactions were incubated for 1 h at 30°C for lamprey and amphioxus Dicer proteins. After the reaction, 500 μL TRIzol was added, RNA was purified using chloroform-isopropanol extraction, and the RNA was resuspended in formamide sample buffer without xylene blue (47.5% formamide, 0.01% SDS, 0.5 mM EDTA). Samples were denatured at 95°C for 5 min and analyzed on a 15% TBE-Urea gel at 400 V for 3 h. Fluorescence imaging was performed using a Bio-Rad imaging system.

### His pull-down assay

LcIFN2 was cloned into the pET-28a vector and expressed in *E. coli* Rosetta cells. LcCRFBs-ECD were cloned into the pcDNA3.1(+) vector with a C-terminal Flag tag. Recombinant LcCRFBs-ECD-Flag proteins were purified from 293F cells. For the pull-down assay, purified His-tagged rLcIFN2 was first incubated with Ni Sepharose 6 Fast Flow (Cytiva) in binding buffer (100 mM Na₂HPO₄, 10 mM NaH₂PO₄, 500 mM NaCl, 20 mM imidazole) for 2 h at 4°C with gentle rotation. The beads were then washed three times with binding buffer, followed by two washes with binding buffer containing 40 mM imidazole to reduce nonspecific binding. Subsequently, purified LcCRFB7-ECD-Flag or LcCRFB14-ECD-Flag proteins were added to the LcIFN2-bound beads and incubated at 4°C for 2 h with rotation. After incubation, the beads were washed five times with binding buffer. The beads were then resuspended with SDS loading buffer, boiled at 100°C for 10 min, and subjected to SDS-PAGE analysis.

### Statistical analysis

Student’s *t*-test was performed to calculate the *p* value between groups by GraphPad Prism (GraphPad Software).

## Data availability

The raw data for scRNA-seq, bulk RNA-seq and small RNA-seq have been deposited at NCBI under the project accession number PRJNA1271237. The codes for genomic screening for *Ifn* like genes are available at https://github.com/SYSUYuanLab/Lamprey.

## Additional materials and methods

Methods for gene cloning and plasmid construction, recombinant proteins purified from cell culture supernatant of HEK293T cells, RNA-seq (Illumina) transcriptome analysis, confocal imaging, plaque assay, Quantitative real-time PCR (qRT-PCR) analysis, FACS analysis, streptavidin-displacement assay, and transposable element annotation and terminal inverted repeat identification are described in the Supplementary Information.

## Acknowledgments

This work was supported by Guangdong S&T Department (2024B1515040009, 2024B1111130003 and 2023B1212060028) and National Natural Science Foundation of China (31970852 and 32170888). The funders had no role in study design, data collection and analysis, decision to publish, or preparation of the manuscript.

## List of Supplementary Materials

Supplementary Materials and Methods

Figure S1. Phylogenetic analysis of lancelet and lamprey IFNs using IL-10 as outgroup.

Figure S2. Establishing the VSV infection model of lamprey.

Figure S3. The CRFBs homologs in lampreys.

Figure S4. Expression pattern of LreCRFBs in various cell types.

Figure S5. Phylogenetic analysis of lamprey IRF and STAT proteins (ML Tree).

Figure S6. LcIFN2 plays broader immunomodulatory roles.

Figure S7. The composition and genomic distribution of the indicated *Ifn* loci.

Dataset S1. List of sequences and referenced accession numbers.

Dataset S2. Abbreviations of species names.

Dataset S3. Sequence alignment of Dicer proteins across species.

Dataset S4. Transposon distribution in the genomic region containing the IFN genes across species.

Dataset S5. List of antibodies used in this study.

Dataset S6. Primers used for gene cloning and qRT-PCR.

## Author contributions

Conceptualization: S.C.Y., A.L.X. and X.L.W.

Methodology: S.C.Y., X.L.W., and Q.H.W.

Investigation: X.L.W., Q.H.W., A.N.X., L.S.D., Z.W.H., Y.G.C., Y.M.Z., R.F., W.C.H., Y.H.C., G.L. and S.C.Y.

Visualization: X.L.W., Q.H.W., and A.N.X.

Writing - original draft: S.C.Y. and X.L.W.

Writing - review and editing: S.C.Y. and A.L.X.

Supervision: S.C.Y. and A.L.X.

Funding acquisition: S.C.Y. and A.L.X.

